# Primary cilia are present on endothelial cells of the hyaloid vasculature but are not required for the development of the blood-retinal barrier

**DOI:** 10.1101/831578

**Authors:** Lana M. Pollock, Brian Perkins, Bela Anand-Apte

## Abstract

Endothelial cilia are found in a variety of tissues including the cranial vasculature of zebrafish embryos. Recently, endothelial cells in the developing mouse retina were reported to also possess primary cilia that are potentially involved in vascular remodeling. Fish carrying mutations in intraflagellar transport (*ift*) genes have disrupted cilia and have been reported to have an increased rate of spontaneous intracranial hemorrhage (ICH), potentially due to disruption of the sonic hedgehog (shh) signaling pathway. However, it remains unknown whether the endothelial cells forming the retinal microvasculature in zebrafish also possess cilia, and whether endothelial cilia are necessary for development and maintenance of the blood-retinal barrier (BRB). In the present study, we found that the endothelial cells lining the zebrafish hyaloid vasculature possess primary cilia during development. To determine whether endothelial cilia are necessary for BRB integrity, *ift57, ift88*, and *ift172* mutants, which lack cilia, were crossed with the double-transgenic zebrafish strain *Tg(l-fabp:DBP-EGFP;flk1:mCherry)*. This strain expresses a vitamin D-binding protein (DBP) fused to enhanced green fluorescent protein (EGFP) as a tracer in the blood plasma, while the endothelial cells forming the vasculature are tagged by mCherry. The Ift mutant fish develop a functional BRB, indicating that endothelial cilia are not necessary for early BRB integrity. Additionally, although treatment of zebrafish larvae with shh inhibitor cyclopamine results in BRB breakdown, the Ift mutant fish were not sensitized to cyclopamine-induced BRB breakdown.

## Introduction

Primary cilia are specialized plasma membrane protrusions that play a role in a number of physiological processes including cell signaling, chemical sensation, and control of cell growth (1). Most nucleated cells in vertebrates possess one immotile primary cilium and perturbation of the structure and function of this organelle leads to abnormal development and organogenesis. Cilia have been observed on endothelial cells both in culture and *in vivo* (2, 3). In developing zebrafish embryos, cilia have been reported on endothelial cells forming the cranial vessels (4) as well as on the caudal artery and vein (5). More recently, endothelial cilia were reported to play a role in the detection of shear stress and vascular remodeling during retinal development (6). The formation and maintenance of primary cilia requires a complex of intraflagellar transport (Ift) proteins (7, 8). Zebrafish embryos with mutations in *ift* genes that lack cilia or have disrupted cilia (9–12) were reported to exhibit an increased incidence of intracranial hemorrhage (ICH), suggesting that endothelial cilia may play a role in maintaining vascular integrity during development (4).

ICH is associated with breakdown of the blood-brain barrier (BBB) (13), and BBB disruption has been demonstrated to precede ICH (14). Similar to the BBB, the endothelial cells lining the retinal microvasculature form the inner blood-retinal barrier (BRB), which serves to protect the neural retinal tissue from the circulating blood. The endothelial cells forming the inner BRB and the BBB are distinct from endothelial cells lining other vascular beds in that they are closely connected by tight junctions, lack fenestrations, and display minimal pinocytotic activity (15).

While the inner BRB is very similar the BBB (16), ultrastructural analysis and comparison of the rat BRB and BBB revealed that the endothelial cells forming the inner BRB have a higher density of junctions and vesicles than those forming the BBB (17). Differences in the permeability of the BRB and BBB to some drugs have also been observed (17, 18). Thus, while the inner BRB and BBB are highly similar, there are strong indications that there may be some unique differences in the structure and functioning of these two barriers

A recent report suggested that cilia on endothelial cells lining the brain vasculature were essential for development of the BBB via the sonic hedgehog signaling pathway (4). Additionally, sonic hedgehog signaling has been shown to be involved in the maintenance of retinal endothelial cell tight junctions in culture (19), suggesting that this signaling pathway could potentially play a role in BRB integrity.

To determine the role of endothelial cilia and hedgehog signaling in BRB integrity *in vivo*, we used the *Tg(l-fabp:DBP-EGFP;flk1:mCherry)* transgenic model of zebrafish BRB development (20). Zebrafish develop a functional BRB by 3 days post-fertilization (dpf) in their hyaloid vessels. We found that the hyaloid vessels of larval zebrafish possess endothelial cilia, but these cilia are not necessary for early integrity of the BRB in IFT mutants.

## Materials and Methods

### Zebrafish

Zebrafish (*Danio rerio*) carrying mutations *ift57^hi3417/curly^*, *ift172^hi2211/moe^* (21), and *ift88^tz288b/oval^* (22), referred to as *ift57, ift172*, and *ift88* herein, were crossed with double-transgenic strain *Tg(l-fabp:DBP-EGFP:flk1:mCherry)* (20, 23, 24). Additionally, transgenic strain *Tg(bactin2:Arl13b-GFP)* was used (25). All animal experimentation was conducted with the approval of the Cleveland Clinic Institutional Animal Care and Use Committee (protocol number 2016-1769).

### Imaging of endothelial cilia

*Tg(flk1:mCherry;bactin2:Arl13b-GFP)* zebrafish embryos and larvae at 2, 3, and 5 dpf were fixed overnight in 4% paraformaldehyde at 4°C. Following washes in phosphate-buffered saline, the lenses and attached hyaloid vessels were isolated as previously described (26). Lenses were placed in coverslip-bottom dishes with Vectashield containing DAPI (Vector Labs) and the hyaloid vasculature was imaged using a Leica SP8 confocal microscope with the 63 x objective.

### Detection of ICH

To inhibit development of pigmentation, zebrafish embryos were raised in embryo medium containing 0.003% 1-phenyl 2-thiourea (PTU). Embryos and larvae were examined for ICH daily by light microscopy.

### Cyclopamine treatment

For early cyclopamine treatment, zebrafish embryos were treated at 1 dpf with 1 μM – 30 μM cyclopamine (Cayman Chemical Company) diluted in embryo medium containing 1% dimethyl sulfoxide (DMSO), and examined for ICH at 2 dpf. For later treatment, zebrafish larvae were treated with 10 μM cyclopamine starting at 5 dpf and assessed for BRB breakdown at 7 dpf.

### Imaging of the BRB

Zebrafish larvae were anesthetized in 0.14 mg/ml ethyl 3-aminobenzoate methanesulfonic acid salt (#118000500, Acros Organics) and examined daily under a Zeiss Axio Zoom V16 fluorescent microscope for leakage of DBP-EGFP from the hyaloid vessels from 3 dpf – 7 dpf.

### Statistics

Prism Software (GraphPad; v5.02) was used for all statistical analyses and graph generation. Results are expressed as means ± SEM. Significance was tested using Fisher’s exact test or t-test.

## Results

### Endothelial cilia are found on the zebrafish hyaloid vasculature

To determine whether the endothelial cells forming the hyaloid vasculature of zebrafish larvae possess primary cilia, the lenses and attached hyaloid vessels were isolated from 2, 3, and 5 dpf *Tg(flk1:mCherry;bactin2:Arl13b-GFP)* zebrafish embryos and larvae and imaged by confocal microscopy (Fig. 1A). The Arl13b protein localizes to the ciliary axoneme, and *Tg(bactin2:Arl13b-GFP)* fish, which stably express an Arl13b fluorescent fusion protein under the ubiquitous β-actin promoter, can be used to label cilia (25, 27). Arl13b-GFP-tagged cilia were present on 53.7 ± 4.9% of hyaloid endothelial cells at 2 dpf, when the hyaloid vessels are first observed at the back of the zebrafish lens (28). However, the number of cilia rapidly decreased to 22.2 ± 6.7% by 3 dpf, and endothelial cilia were not observed at 5 dpf (Fig. 1B). This observation is similar to observations in the zebrafish caudal aorta and caudal vein, in which 76% of ECs possess cilia during vascular morphogenesis, while only 4% of ECs possess cilia during later developmental stages (5).

**Fig. 1.**
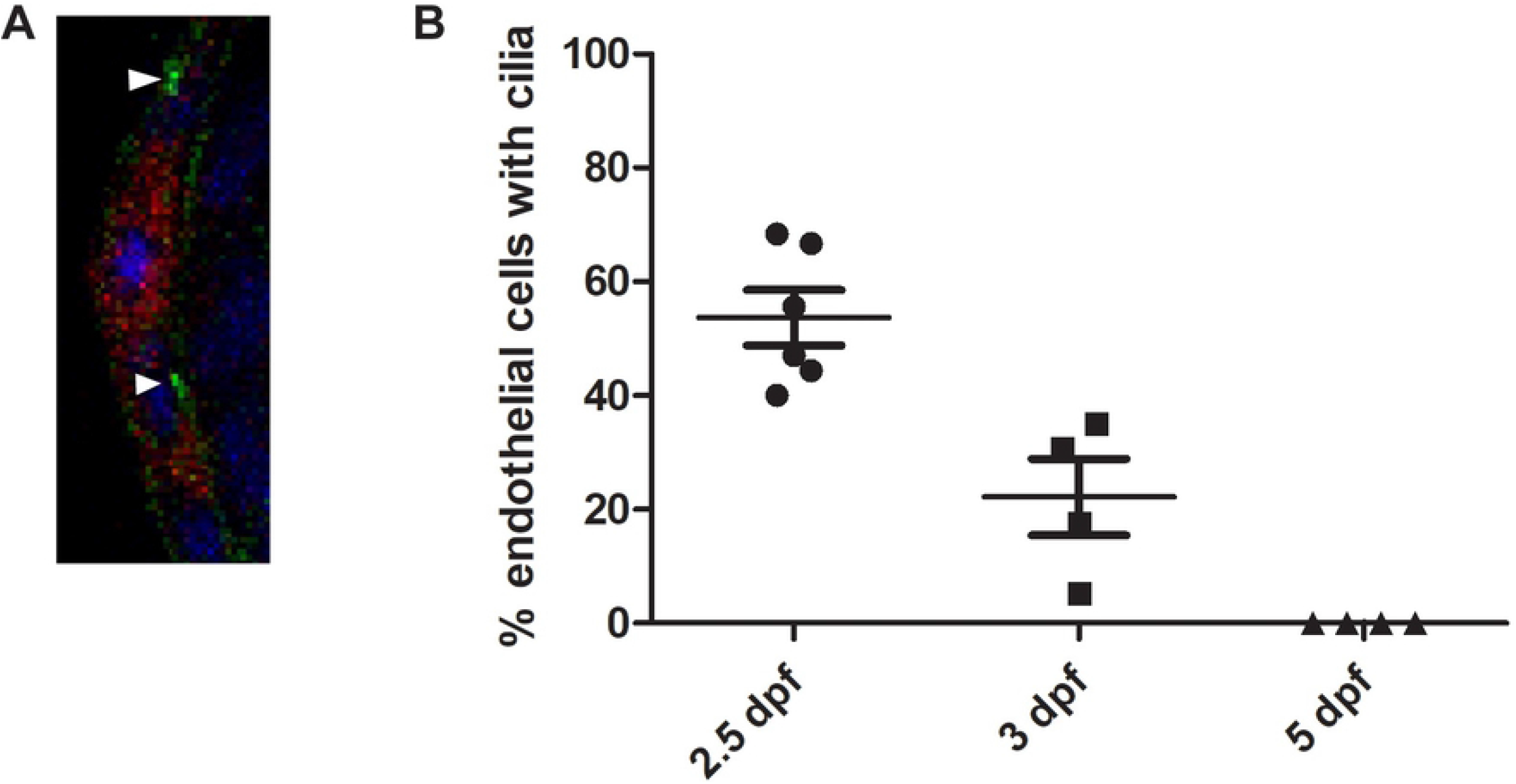
Endothelial cilia are observed during early development of the hyaloid vasculature, but rapidly decline in number by 3 dpf. A) Confocal micrographs of the hyaloid vessels (red) of a 3 dpf *Tg(flk1:mCherry;bactin2:Arl13b-GFP)* embryo. Cilia (green, arrowheads) were observed on the endothelial cells, near the nuclei (DAPI, blue). B) Percentage of hyaloid vessel endothelial cells possessing a primary cilium at 2, 3, and 5 dpf (n > 80 endothelial cells from ≥ 4 eyes per time point). Error bars = SEM.

### Endothelial cilia are not necessary for development of the blood-retinal barrier

To determine whether endothelial cilia are necessary for BRB development, we evaluated the BRB in live *Tg(l-fabp:DBP-EGFP;flk1:mCherry);ift172, Tg(l-fabp:DBP-EGFP;flk1:mCherry);ift57*, and *Tg(l-fabp:DBP-EGFP;flk1:mCherry);ift88* larvae. These IFT mutants fail to form cilia (9, 29). IFT homozygous mutant larvae were initially identified by screening for the previously-described characteristic ventral body curvature and kidney cyst formation phenotypes (21), and later confirmed by genotyping. We have previously demonstrated that zebrafish develop a fully functional BRB by 3 dpf (IFT mutant larvae all had an intact BRB at 3 - 6 dpf, indicated by localization of the DBP-EGFP endogenous blood plasma tracer within the mCherry-tagged vessels (Fig. 2, n = 50 fish per group). Thus, endothelial cilia do not appear to be necessary for the development and early maintenance of the BRB. Most IFT mutant larvae died at around 9 dpf, precluding analysis at later developmental stages. As a positive control for BRB breakdown in the *Tg(l-fabp:DBP-EGFP;flk1:mCherry)* model, transgenic non-mutant embryos were treated with 0.15 μM BMS493, an inhibitor of retinoic acid signaling previously demonstrated to disrupt the BRB (30), from 2 dpf until 7 dpf (Fig. 2E).

**Fig. 2.**
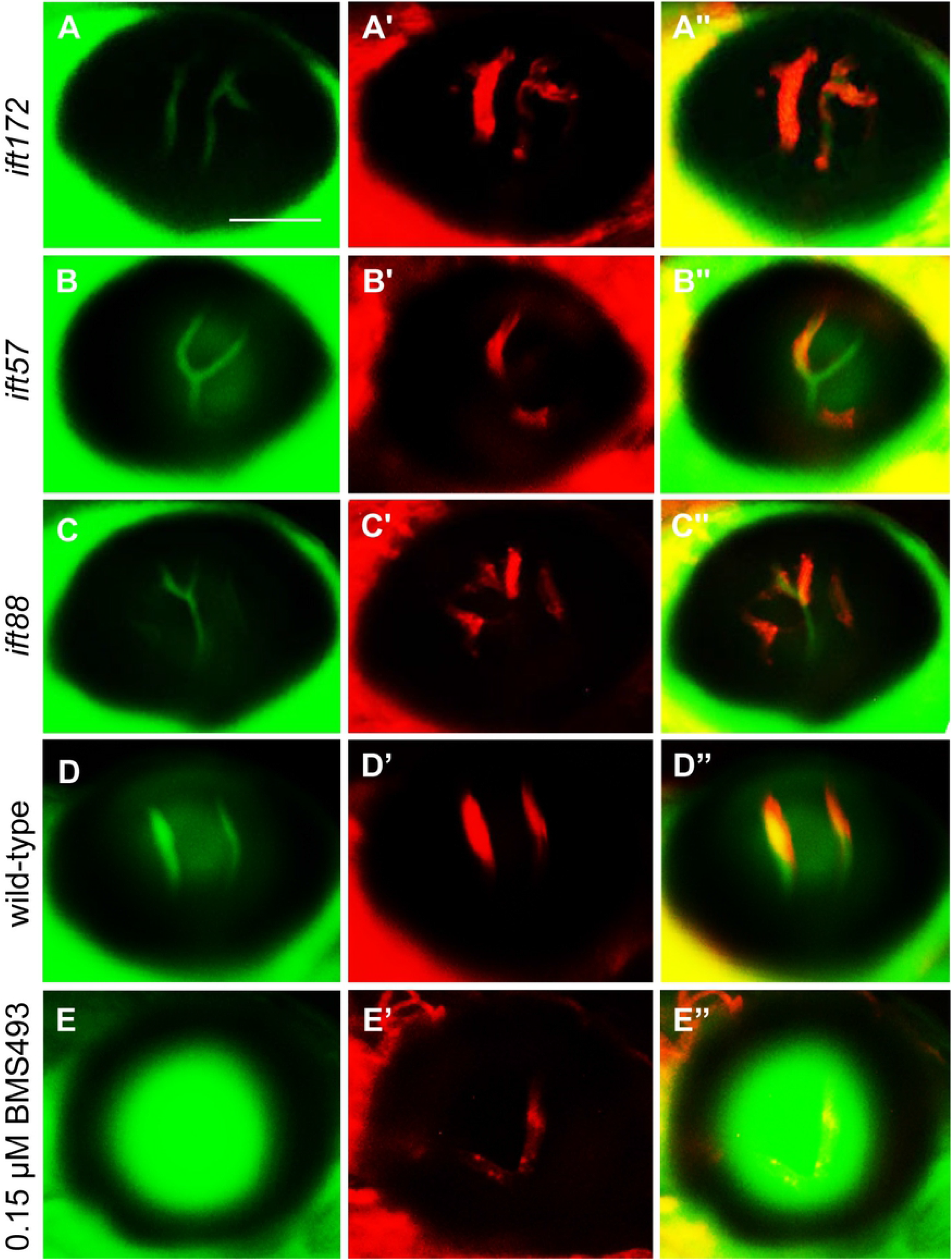
IFT mutant fish develop and maintain a functional BRB. Fluorescent micrographs of the hyaloid vessels in live 6 dpf IFT mutant and wild-type *Tg(l-fabp:DBP-EGFP;flk1:mCherry)* fish. In the *ift172* (A), *ift57* (B), and *ift88* (C) mutants and wild-type larvae (D), the DBP-EGFP-associated signal (green) localizes within the mCherry-tagged vessels (red), indicating that the BRB is intact. (E) As an example of BRB breakdown, *Tg(l-fabp:DBP-EGFP;flk1:mCherry)* embryos were treated with 0.15 μM BMS493, an inhibitor of retinoic acid signaling, from 2 dpf until 7 dpf. Scale bars = 50 μm

### Low rates of intracranial hemorrhage in zebrafish and *ift172*, *ift57*, *ift88* intraflagellar transport mutants

Kallakuri *et al*. (4) previously reported an ICH rate of 34% at starting at 2 dpf in *ift172* mutants, although this phenotype was not fully penetrant, with variable frequencies of ICH observed between different families. Another study by Eisa-Beygi *et al*. (31) reported an ICH rate of 11.8% at a similar developmental timepoint in *ift172* mutant larvae. To determine whether the families used in our study also displayed an increased rate of ICH, embryos were treated with 0.003% PTU to reduce pigmentation and examined for cranial hemorrhages each day from 2-4 dpf. The highest ICH rates observed among all families screened were 12.5%, 15.3%, and 16.6% in the *ift172*, *ift57*, and *ift88* mutants, respectively (Fig. 3). While these rates are significantly higher than the 0 - 3% ICH rates observed in their wild type and heterozygous siblings, the majority of clutches had 0% ICH in the IFT mutant embryos (15/18 *ift172* clutches, 13/15 *ift57* clutches, and 6/10 *ift88* clutches screened). Additionally, most incidences of ICH were not detected until 3 dpf, and usually resolved by 4 dpf (Fig. S1).

**Fig. 3.**
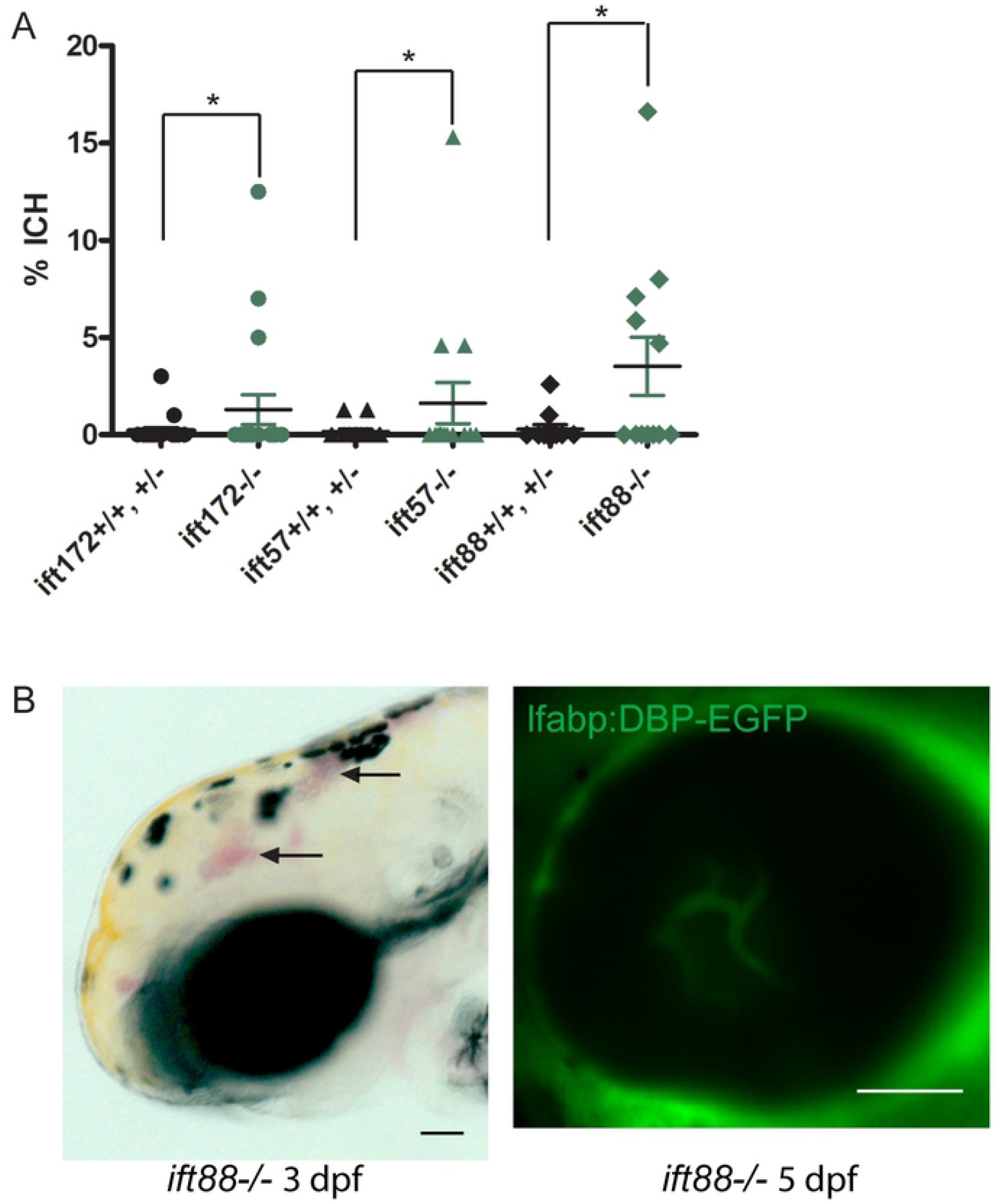
*Ift* mutant fish with ICH maintain an intact BRB. Embryos were screened daily for ICH and BRB integrity was assessed at 5 dpf. (A) Average percentages of *ift172^-/-^*, *ift57^-/-^*, and *ift88^-/-^* embryos and their wild type and heterozygous siblings with ICH (n = 18 *ift172* crosses, 15 *ift57* crosses, and 6 *ift88* crosses of ≥ 50 embryos each). Error bars = SEM. *P < 0.05 by Fisher’s exact test. (B) An *ift88^-/-^* embryo with ICH at 3 dpf (bright field image at left, arrowheads) and an intact BRB at 5 dpf (confocal micrograph at right). Scale bar = 50 μm.

### Inhibition of hedgehog signaling does not sensitize zebrafish *ift57, ift88*, and *ift172* mutants to BRB breakdown

In the previous report by Kallakuri et al., *ift81-*mutant fish were sensitized to ICH induced by treatment with cyclopamine, an antagonist of the hedgehog signaling pathway, even in families without spontaneous ICH. To determine whether the IFT mutant fish in this study were likewise sensitized to cyclopamine-induced ICH, fish were subjected to cyclopamine treatment following the protocol used by Kallakuri et al. in which embryos were treated from 25 hours post-fertilization (hpf), and analyzed for ICH at 52 hpf. We initially treated IFT mutant embryos and their siblings with the 30 μM concentration used by Kallakuri et al. As previously reported, this concentration did not significantly affect embryo morphology. However, treatment with the 30 μM concentration resulted in severe ICH in 98-100% of all treated fish, making it difficult to determine whether the IFT mutant fish were sensitized. High variability in the response of different zebrafish strains to cyclopamine treatment has been previously reported (32). Thus, we titrated the cyclopamine dosage to look for a concentration at which the mutants are sensitized compared to their wild type and heterozygous siblings. Nearly 100% of the treated fish developed ICH in concentrations as low as 5 μM. Treatment with 1 μM cyclopamine resulted in ICH in 20-30% of all treated fish, though the incidence of ICH was not significantly different between the IFT mutant fish and their wild-type and heterozygous siblings (Fig. 4A).

**Fig. 4.**
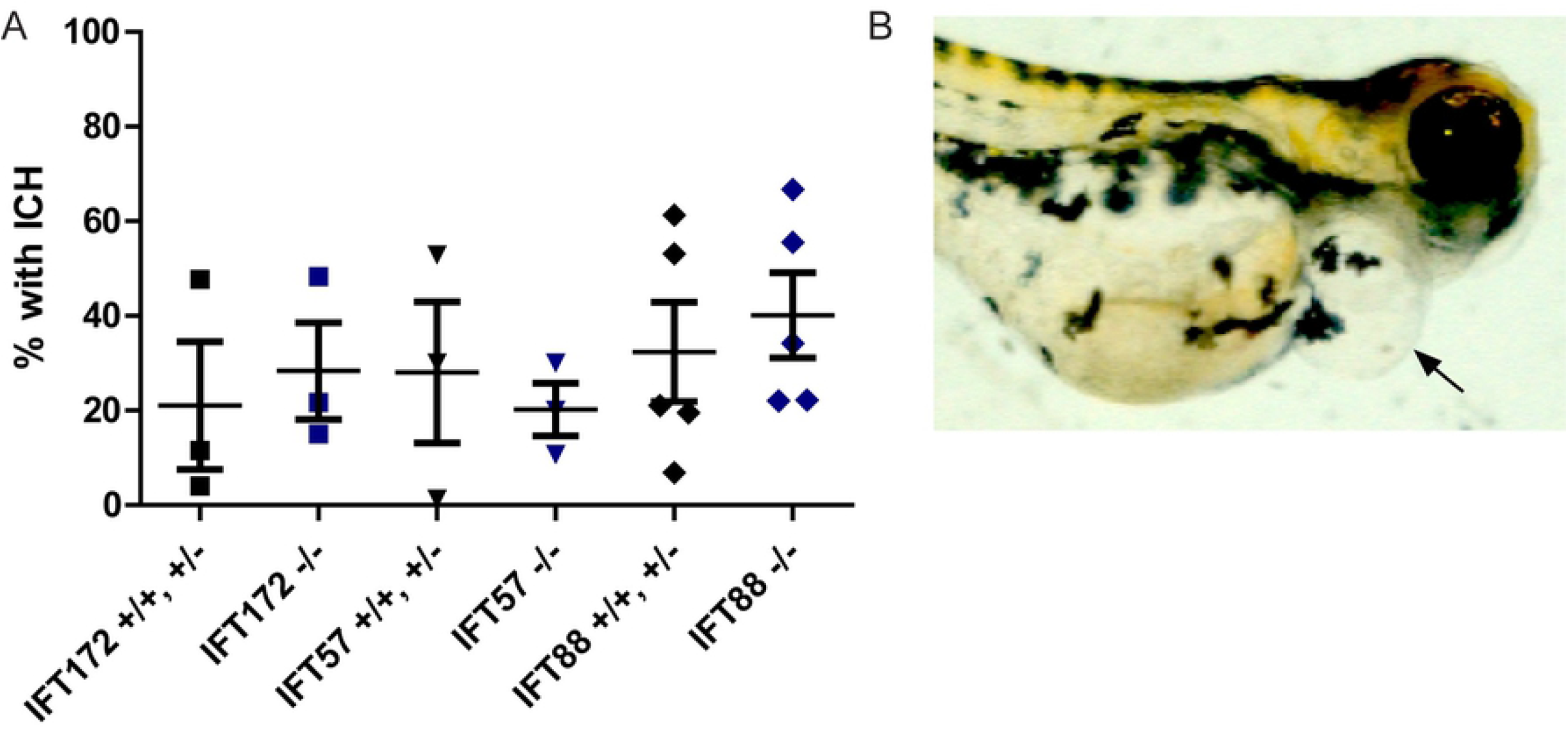
Early inhibition of the hedgehog signaling pathway results in severe edema and BRB non-perfusion. Zebrafish embryos were treated with 1 μM cyclopamine starting at 25 hpf. (A) Percentages of cyclopamine-treated IFT mutant and non-mutant embryos that developed ICH by 52 hpf. (B) Image of a 6 dpf wild-type fish that had cyclopamine-induced pericardial edema (arrow).

Treatment with 1 μM cyclopamine starting at 25 hpf did not affect the BRB, although it resulted in pericardial edema in 97.3% of treated wild type zebrafish larvae by 6 dpf (Fig. 4B). While treatment with 10 μM cyclopamine starting later in development (5 dpf) rarely resulted in edema/non-perfusion of the hyaloid vessels (Fig. 5B) in wild-type larvae, the number of fish with edema/non-perfusion was variable in the clutches of *ift* mutant larvae and their wild-type and heterozygous siblings (Fig. 5C). Among the clutches without edema/non-perfusion, 10 μM cyclopamine treatment starting at 5 dpf resulted in BRB breakdown in 82.8 ± 8% of wild-type larvae by 7 dpf (Fig. 5A). No significant difference in the rate of cyclopamine-induced BRB breakdown was observed in the IFT mutant larvae (Fig. 5C). This indicates that hedgehog signaling is necessary for BRB integrity, although endothelial cilia are likely not involved in this maintenance.

**Fig. 5.**
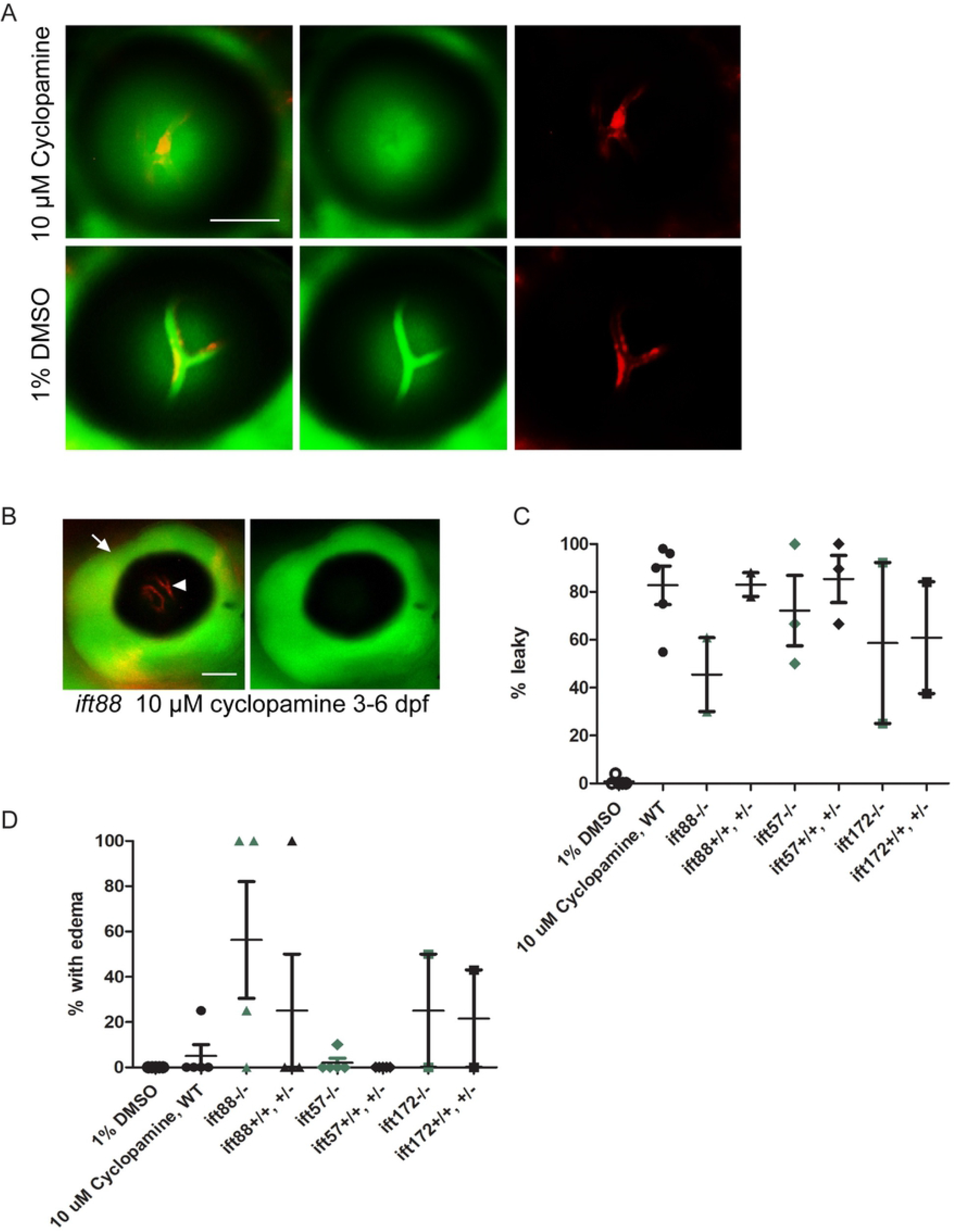
Later treatment of zebrafish larvae with higher doses of cyclopamine results in BRB leakage. Zebrafish larvae were treated with 10 μM cyclopamine from 5 dpf – 7 dpf. (A) Cyclopamine-treated larvae had BRB breakdown at 7 dpf, as seen by the detection of DBP-EGFP (green) outside of the hyaloid vessels (red). (B) Fluorescent micrograph of a cyclopamine-treated zebrafish at 7 dpf with edema around the eye (arrow), and non-perfusion of the hyaloid vasculature (arrowhead). (C) Graph depicting the percentage of WT and Ift mutant fish in each clutch with edema/non-perfusion after treatment with 10 μM cyclopamine from 5 dpf – 7 dpf. (D) Graph depicting the percentage of WT and Ift mutant fish in clutches without edema/non-perfusion that had BRB breakdown upon treatment with 10 μM cyclopamine or 1% DMSO vehicle control. Each data point in (C) and (D) represents a clutch of >50 larvae. Error bars represent SEM. Scale bars = 50 μm.

## Discussion

Endothelial cilia have been proposed to serve a number of different roles including hedgehog signaling (4), mechanosensation of blood flow (33), and nitric oxide signaling (34). We found that the endothelial cells lining the inner retinal vasculature possess cilia during BRB development, though the number of hyaloid endothelial cells possessing a primary cilium rapidly declines by 3 dpf. It is possible that this decline is due to loss of activity of the *actb2* promoter driving Arl13b-GFP expression. However, a similar decrease in the number of endothelial cells possessing a primary cilium was reported in the arteries of the developing mouse retina and in the zebrafish caudal aorta (5, 6).

Endothelial cilia do not appear to be necessary for development of hyaloid vascular integrity. It is possible that endothelial cilia may be necessary for long-term maintenance of the BRB at adult stages. However, we were unable to assess this due to mortality of the IFT mutant fish at larval stages. Knockout of IFT88 in the mouse endothelial cells has been reported to result in transient decreased radial expansion, decreased vascular density, and fewer sprouting cells at the front of the plexus in the retina (6). Changes were not observed in the developing hyaloid vasculature of *ift57, ift172*, and *ift88* mutant zebrafish, though this may be due to differences in the developmental processes involved in the mouse and zebrafish retinal vasculature.

While higher rates of ICH have been reported in IFT mutant zebrafish (4), developmental ICH was not detected in the majority of clutches screened in this study. Also contrary to the previous report, our IFT mutant fish did not appear to be more sensitized to cyclopamine treatment-induced ICH than their wild-type and heterozygous siblings. These contrasting results may be due to other currently unknown genetic risk factors that vary between the fish stocks used in the respective studies.

Sonic hedgehog signaling has been proposed to be involved in establishing vascular integrity in both the brain and the retina (4, 19). While we did not observe an increased incidence of ICH in IFT mutant fish upon treatment with cyclopamine at 25 hpf, later treatment at 5 dpf with higher doses of cyclopamine resulted in BRB breakdown. No differences in the rates of cyclopamine-induced BRB breakdown were detected between IFT mutant larvae and their non-mutant siblings. This indicates that hedgehog signaling is necessary for BRB integrity, though endothelial cilia are not necessary for this process.

## Acknowledgements

Thanks to Mariya Ali and Alecia Cutler for technical assistance. We also thank Drs. Didier Stainier, Suk-Won Jin, and Neil Chi for providing the *Tg(flk1:mCherry)* fish. This work was supported by NIH grants EY016490, EY015638, EY026181, EY017037, T32EY024236, P30EY025585, and a Research to Prevent Blindness Challenge Grant; a Doris and Jules Stein Professorship Award from Research to Prevent Blindness, and a Foundation Fighting Blindness Center Grant.

---

**Supplemental Figure 1: ICH develops in some *ift* mutant zebrafish larvae at 3 dpf and resolves by 4 dpf.** Bright-field images of representative *ift172* (A), *ift57* (B), and *ift88* (C) fish with ICH (arrowheads) at 3 dpf. The same fish were imaged again the following day (4 dpf), and the areas of ICH were largely resolved (D-F, arrowheads).

